# Occurrence, identification and antibiogram signatures of selected Enterobacteriaceae from Tsomo and Tyhume rivers in the Eastern Cape Province, Republic of South Africa

**DOI:** 10.1101/2020.08.11.246025

**Authors:** Folake Temitope Fadare, Martins Ajibade Adefisoye, Anthony Ifeanyi Okoh

**Affiliations:** SAMRC Microbial Water Quality Monitoring Centre, University of Fort Hare, Alice, South Africa; Applied and Environmental Microbiology Research Group, Department of Biochemistry and Microbiology, University of Fort Hare, Alice, South Africa

**Keywords:** Multidrug-resistant, Enterobacteriaceae, Extended-spectrum β-lactamase, plasmid-mediated AmpC

## Abstract

The increasing occurrence of multidrug-resistant Enterobacteriaceae in clinical and environmental settings has been seen globally as a complex public health challenge, mostly in the developing nations where they significantly impact on freshwater used for a variety of domestic purposes and irrigation. In this paper, we give details as regards the occurrence and antibiogram signatures of Enterobacteriaceae community in Tsomo and Tyhume rivers within the Eastern Cape Province, Republic of South Africa using standard methods. Average distribution of the presumptive Enterobacteriaceae in the rivers ranged from 1 × 10^2^ CFU/100ml to 1.95 × 10^4^ CFU/100ml. We confirmed 56 (70.8%) out of 79 presumptive Enterobacteriaceae isolated being species within the family Enterobacteriaceae through the Matrix-Assisted Laser Desorption Ionization Time of Flight technique. From this, *Citrobacter*-, *Enterobacter*-, *Klebsiella* species and *Escherichia coli* were selected (n=40) due to their pathogenic potentials for antibiogram profiling. The results of the antibiotic susceptibility testing gave a revelation that all the isolates were phenotypically multidrug-resistant while resistance against ampicillin (95%), tetracycline and doxycycline (88%) and trimethoprim-sulfamethoxazole (85%) antibiotics were most prevalent. The Multiple Antibiotic Resistance indices stretched from 0.22 to 0.94, with the highest index observed in a *C. freundii* isolate. Molecular characterisation using the PCR technique revealed the dominance of *bla*_TEM_ (30%; 12/40) among the ten groups of extended-spectrum β-lactamase (ESBL) genes assayed. The prevalence of others was *bla*_CTX-M_ genes including group 1, 2 and 9 (27.5%), *bla*_SHV_ (20%), *bla*_KPC_ (17.6%), *bla*_GES_ (11.8%), *bla*_IMP_ (11.8%), *bla*_VIM_ (11.8%), *bla*_OXA-1-like_ (10%), *bla*_PER_ (2.5%), *bla*_OXA-48-like_ (5.9%) and *bla*_VEB_ (0%). From the six plasmid-mediated AmpC (pAmpC) genes investigated *bla*_ACC_, *bla*_EBC_, *bla*_FOX_, *bla*_CIT_, *bla*_DHA_ and *bla*_MOX_, only the first four were detected. In this category, the most dominant was *bla*_EBC,_ with 18.4% (7/38). The prevalence of the non-β-lactamases include *tetA* (33.3%), *tetB* (30.5%)*, tetC* (2.8%), *tetD* (11.1%), *tetK* (0%), *tetM* (13.9%), *catI* (12%), *catII* (68%), *sulI* (14.3%), *sulII* (22.8%) and *aadA* (8.3%). Notably, a *C. koseri* harboured 42.8% (12/28) of the genes assayed for which includes five of the ESBL genes (including the only *bla*_PER_ detected in this study), two of the pAmpC resistance genes (*bla*_ACC_ and *bla*_CIT_) and five of the non-β-lactamase genes. To the best of our knowledge, this study gives the first report on *C. koseri* exhibiting co-occurrence of ESBL/AmpC β-lactamase genes from the environment. The detection of a *bla*_PER_ producing *Citrobacter* spp. in this study is remarkable. These findings provide evidence that freshwater serves as reservoirs of antimicrobial resistance determinants which can then be easily transferred to human beings via the food chain and water.

## Introduction

Globally, the appearance, widespread and distribution of antimicrobial resistance in bacteria have been described as a complex public health challenge (1–3). Initially, it was perceived as a problem restricted to clinical settings (4). However, findings have shown that genes encoding various antibiotic resistance within the environment predate the discovery of antibiotics used in clinical settings. This discovery, therefore, implicates the environment as the origin of antimicrobial resistance evolution (5–8). In different regions of the world, there have been various studies which have carried out investigation into the occurrence of antibiotic-resistant bacteria (ARB) in various aquatic milieu (5,9–12). Since the first discovery of antibiotics over eight decades ago, they have been used to save millions of lives in treating bacterial infections and diseases. This feat was, however, short-lived as the numbers of organisms becoming resistant to antibiotics have grown at alarming rates than the anticipated normal evolutionary process for microorganisms (2,4,13). The ARB could be selected under certain circumstances such as when soil organisms which naturally can produce antibiotics are deposited to freshwater sources via runoffs. These natural stockpiles of ARB alongside the antibiotic-resistant genes (ARGs) conferring resistance on them could then be a wellspring of transferable traits for emerging pathogens.

Once individual organisms in a particular niche have developed or acquired resistance, they quickly transfer this to other organisms in widely varied niches. Various horizontal gene transfer mechanisms often facilitate this widespread leading to the spread of resistant organisms (13–15). Selective pressure that favours the evolution of resistance occurs whenever antimicrobials are utilised in animal husbandry and aquaculture, as therapeutics, prophylaxis, metaphylaxis and as growth promoters (16–20). This is primarily because by nature antibiotics act on the principle of selective toxicity wherein they kill or inhibit microbial pathogens while little or no damage is done to the host(2,5,16).

The gastrointestinal tract of animals, as well as humans, may serve as an alternate host or passive carrier of ARB, and these can cause different diseases via diverse mechanisms. When antibiotics are applied, ARB survives and multiply rapidly reaching high densities in the intestinal lumen from where they are then excreted into the environment making the possibility of containing their spread far-fetched (21). The ARB may initially be commensals living in the gastrointestinal tract, but after a while may acquire genetic materials such as plasmids, integrons, or transposons carrying various resistance genes and virulence factors, which can then transform these initially harmless bacteria into more virulent and resistant organisms (4,16). Also, a large portion of antibiotics used is not broken down into inactive constituents thereby retaining their properties even after discharge from the body into the environment (4,18,22,23). These sometimes travel to wastewater treatment plants (WWTPs) via domestic sewer lines or are excreted directly into the soil or surface waters, thereby further adding to the antibiotic resistance pressure impacted on the environment (24,25). Surface waters have been reported to be “hotspots” of antimicrobial resistance contamination owing to being a recipient of discharges from diverse sources which include contaminants from industrial, agricultural and domestic settings of varying chemical and microbial concerns (5,18,26,27). Furthermore, the surface water being a reservoir of ARB and ARGs could serve as a dissemination port for the proliferation of new resistant strains which can be enhanced by the acquisition bacteriophages or integrons through horizontal gene transfer mechanisms (28,29).

One of the most prestigious groups of microorganisms that have been popularly implicated in the spread of ARGs in the environment is the Enterobacteriaceae group. Members of this family are among the natural microflora in the gastrointestinal tracts of warmblooded animals including humans and commonly found in diverse environmental sources such as water, plants and soil (16). Clinically significant genera of this family that commonly cause infections are *Citrobacter*-, *Enterobacter*-, *Escherichia coli*, *Klebsiella*-, *Morganella*-, *Plesiomonas*-, *Proteus*-, *Providencia*-, *Salmonella*-, *Serratia*-, *Shigella*- and *Yersinia* species. (16,30,31). They have been commonly implicated in various infections including pneumonia, enteritis, diarrhoea, septicaemia, wound and infections involving the central nervous system (16).

Various ARGs against various classes of antibiotics such as aminoglycosides, tetracyclines, sulphonamides, trimethoprims, β-lactams and carbapenems have been discovered in various aquatic environments (7,9,10,12). The principal mechanism of antibiotic resistance in members of the family Enterobacteriaceae is the production of enzymes called β-lactamases. These enzymes include the Extended-Spectrum β-lactamases (ESBLs) and the plasmid-mediated AmpC-producing (pAmpC) (32). The β-lactams have a particular chemical structure called the β-lactam ring and are the largest and most generally used class of antibiotics. These are mostly semi-synthetic compounds, which emanate from bacteria and fungi within the environment (33,34). They function by preventing the transpeptidation of the peptidoglycan cell wall of the invading bacteria which is necessary for the completion of the cell wall(33–35). Resistance against the β-lactams is principally by the structural modification of the penicillin-binding proteins thereby resulting in decreased attraction of the drugs or worse still, the bacteria produce enzymes, which break the β-lactam ring thereby rendering the drugs ineffective. Other resistance mechanisms include decreased permeability or active transportation of the antibiotics out of the bacterial cell using efflux pumps (35).

The ability of Enterobacteriaceae species to produce enzymes ESBL and pAmpC is a severe emerging problem and a source of concern for environmental safety (1,36–38). These enzymes endow bacteria with various resistances against most β-lactam antibiotics including penicillins, carbapenems and cephalosporins (16,34,39). This ability to produce the ESBL/pAmpC enzymes which cleaves the β-lactam ring thereby conferring resistance on the organism makes it one of the most potent ways of fostering resistance within the Enterobacteriaceae family (38). This scenerio thereby undermines the effectiveness of current antibiotics and unfortunately further hampers the development of new drugs. Initially, carbapenems were prescribed as a last resort for the therapy of gram-negative severe nosocomial infections which are caused by ESBLs and pAmpC producing Enterobacteriaceae. Unfortunately, due to the widespread use of this class of antibiotics, various carbapenem-resistant Enterobacteriaceae has been reported (40,41). The ARB can be readily spread directly to humans when contaminated water are consumed or indirectly via the consumption of foods irrigated with water from these contaminated freshwater sources. This further enables the spread and persistence of ARB and ARGs within our environment and among the general populace (42,43). It is therefore imperative that studies be carried out to monitor the prevalence of ARB and ARGs in freshwater sources. Here, we describe the occurrence and antibiogram signatures of Enterobacteriaceae community in Tsomo and Tyhume rivers in ECP, RSA.

## Materials and methods

### Study sites

Two freshwater resources sampled in this study are the Tyhume River and Tsomo River. The Tyhume River is found in Amathole District Municipality (ADM) while the Tsomo River is located in Chris Hani District Municipality (CHDM) both in the ECP. Within this province, ADM is located in the central part, while CHDM is found in the north-eastern part. They are both mainly involved in agricultural production and serve as a hub for various agricultural processes which may be in part due to their nearness to East London and Port Elizabeth seaports.

The Tyhume River is situated in the Raymond Mhlaba local municipality within ADM. The river originates from the mountains in Hogsback and passes along several rural settlements along its path to Alice and finally empties to the Kieskamma River at Manqulweni community. This river is used by the different host communities it passes through for diverse domestic activities such as washing, drinking, cooking, recreational activities and religious activities which include baptism especially in locations where potable water is not within reach. It is also channelled into the Binfield Park Dam, which is a hub for the supply of raw water to the different water treatment plants in that vicinity. The treated water is then subsequently redistributed to Alice town and the communities in that area as a source of potable water. The four sampling sites along the Tyhume River include Hala, Khayalethu, Sinakanaka and Alice.

The Tsomo River is found in the Intsika Yethu local municipality within CHDM. This river originates about 10 km to the northwest of the town Elliot and flows through several rural settlements including Tsomo, Cala, and Ncora. It flows southwards and finally empties into the Great Kei River. This river is also utilised by the different host communities it passes through for various domestic purposes. The river is also channelled into the Ncora Dam, which is used solely for irrigation purposes. The details of these sample collection points are described in supplementary Table 1.

### Sample collection and processing

In October and November 2017, water samples were collected from four and two points on the Tyhume and Tsomo River, respectively in triplicates on a once-off basis. Using sterile 1 L polypropylene bottles samples were retrieved from each site in triplicates. These were maintained at 4 °C and immediately transported to the laboratory and analysed within 6 hours. Ten-fold serial dilution (10^−1^, 10^−2^ and 10^−3^) was carried out, then 100 mL of the appropriately diluted sample was filtered through a cellulose nitrate membrane filter with a pore size of 0.45 μm (Sartorius, Goettingen, Germany) using a vacuum pump. All samples were processed in triplicates. After each filtration, the membrane filter was aseptically placed onto Petri-dishes containing sterile Violet Red Bile Glucose (VRBG) agar (Conda – pronadisa, USA) plates and incubated at 37 °C for 24 hours for the enumeration and isolation of presumptive Enterobacteriaceae. After incubation, purple-red colonies with a diameter of at least 0.5 mm were counted as characteristic presumptive Enterobacteriaceae. The results obtained were recorded as colony-forming units per 100 ml (CFU /100ml) (44).

### Isolation and MALDI-TOF identification of presumptive targeted Enterobacteriaceae

The isolation was executed following standard procedures utilising general and selective media. Morphologically different colonies were sub-cultured on sterile Eosin Methylene Blue differential agar (EMB) (Merck, Darmstadt, Germany) and incubated at 37 °C for 24 hours. Green metallic sheen colonies and pink to dark purple colonies were selected as lactose fermenters (presumptive *E. coli, Citrobacter-, Enterobacter-* and *Klebsiella* species) and were purified using nutrient agar. Pure presumptive isolates were preserved in 20% glycerol stock solutions at −80 °C for future assays. The confirmation of the identification of isolates was obtained by using matrix-assisted laser desorption and ionisation time-of-flight coupled with mass spectrometry (MALDI-TOF MS). The MALDI Biotyper 3.0 software (Bruker Daltonics, Bremen, Germany) was used for the confirmation of the identities of the presumptive organisms to species level. The instrument was calibrated with bacterial test standard (BTS) and all analysis was carried out in duplicates. All results were interpreted following manufacturer’s instructions and results with values less than 1.700 were excluded as they identification were not reliable as described by (45).

### Antibiotic susceptibility testing

The antimicrobial susceptibility patterns of all targeted identified Enterobacteriaceae isolates totalling 40 recovered from the rivers were analysed using the Kirby-Bauer disk diffusion technique following the Clinical and Laboratory Standards Institute guidelines (46). The isolates were subjected to 18 antibiotic discs which belong to 11 different classes of antibiotics which are usually used for treating infections triggered by Enterobacteriaceae as suggested by the Center for Disease Control and Prevention (CDC). The antibiotic classes, the specific antibiotics used along with their concentrations are as follows; Aminoglycosides: gentamicin (GM; 10 μg) and amikacin (AK; 30 μg); β-lactams: amoxicillin/clavulanic acid (AUG; 30 μg) and ampicillin (AP; 10 μg); Carbapenems: imipenem (IMI; 10 μg) and meropenem (MEM; 10 μg); Cephems: cefotaxime (CTX; 30 μg) and cefuroxime (CXM; 30 μg); Fluoroquinolones: ciprofloxacin (CIP; 5 μg) and norfloxacin (NOR: 30 μg); Nitrofurans: nitrofurantoin (NI; 300 μg); Phenicols: chloramphenicol (C; 30 μg); Polymyxins: polymyxin B (PB; 300 units) and colistin sulphate (CO; 25 μg); Quinolones: nalidixic acid (NA; 30 μg); Sulfonamides: trimethoprim-sulfamethoxazole (TS; 25 μg); Tetracyclines: tetracycline (T; 30 μg) and doxycycline (DXT; 30 μg). Briefly, the inoculum was prepared in normal saline solution by dispersing a single colony picked with a sterile cotton swab. The turbidity of the resulting solution was compared to 0.5 McFarland turbidity standards (equivalent to 1.5 × 10^8^) and where necessary, adjusted using sterile normal saline. Then, 100 μl of each of the standardised bacterial test suspension was spread on the Mueller-Hinton agar (Merck, Johannesburg) plates using sterile cotton swabs. Using a disc dispensing apparatus (Mast Diagnostics, U.K),the relevant antibiotic disks (Mast Diagnostics, U.K) were placed at an equidistance of 30 mm on the inoculated plates. The plates were then inverted fifteen minutes after the discs were applied and incubated at 37 °C for 18 – 24 hours. The zones of inhibition were determined by measuring the diameter of the clear zones around the antibiotic disks to the nearest millimetres. The results for the isolates were interpreted as “Resistant (R), Intermediate (I), or Susceptible (S)” following the (46) breakpoints while the class Polymyxins were interpreted using the zone diameter of *E. coli* ATCC 25922.

### Analysis of Multiple Antibiotic Resistance Phenotype (MARP) and Multiple Antibiotic Resistance Index (MARI) of bacterial isolates

The MARP patterns for isolates that displayed resistance against more than three antibiotics out of the tested eighteen were assessed (47). Isolates that were resistant against at least three antimicrobial classes were classified as multidrug-resistant (MDR). The multidrug resistance pattern, the number of antibiotics the isolates displayed resistance against, as well as the number of the phenotypic patterns observed were analysed. The MARI of the individual multidrug isolate was derived using a Mathematical equation as described by Krumperman (1983). The mathematical expression is given as:

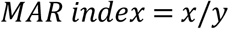

Where “x” is the number of antibiotics against which a bacterial species exhibited resistance, “y” is the total number of antibiotics which that bacterial species was exposed. A MAR index value more than 0.2 is an indicator of intensive usage of antibiotics in that area and predict high-risk vicinity for the possible promotion of antibiotic resistance (48,49).

### Antibiotic resistance genes detection

Purified bacteria isolates were revived by inoculating into a nutrient broth and then incubated overnight at 37 °C. The cultures were then transferred unto nutrient agar (NA) and incubated overnight at 37 °C. The extraction of the total genomic DNA was done through boiling method as reported by (50) with slight modifications. Briefly, with the aid of a sterilised inoculating loop, single colonies from NA were suspended into 100 μl of sterilised nuclease-free water (Thermo Scientific, USA) in sterile Eppendorf tubes (Biologix, USA). The resulting suspension was swirled with a vortex mixer (Digisystem Laboratory, Taiwan). Boiling at 100 °C for 10 minutes with an AccuBlock (Digital dry bath, Labnet) lysed the cells. These were incubated immediately on ice for 5 mins after that the cell debris was extracted by centrifuging at 13,500 rpm for 10 minutes using a Mini-spin micro-centrifuge (LASEC, RSA). Cell lysate supernatant was removed into sterile Eppendorf tubes which were utilised as template DNA in PCR assays straight away or stowed at −20 °C for future assays. Different resistance determinants were assayed for in the targeted bacterial isolates showing full or intermediate resistance. Nineteen resistance genes that encode β-lactamases, various variants of ESBL and pAmpC resistance gene determinants were assayed for in duplex and multiplex PCR protocols as reported by (51). Each gene with the primer set used as well as their expected molecular sizes is described in Supplementary Table 2. The thermal cycling conditions for the PCR assays were as follows: initial denaturation at 94 °C for 10 mins; followed by 30 cycles of 94 °C for 40s, 60 °C for 40s, and 72 °C for 1 min; and then a final elongation step at 72 °C for 7 mins. For the carbapenemase genes (*bla*_VIM_, *bla*_IMP_ and *bla*_KPC_) and (*bla*_GES_ and *bla*_oxa-48_) multiplex PCR assays, the annealing temperature was optimal at 55 °C and 57 °C respectively. Singleplex, duplex and multiplex PCR protocols as reported by (9) were employed for the detection of 12 resistance genes that code for resistance against the non-β-lactams which include the sulphonamides, tetracyclines, phenicols and aminoglycosides. Supplementary Table 3 shows the list of the genes assayed with their expected molecular sizes.

For the PCR assays, a Mycycler™ Thermal Cycler PCR system (BioRad, USA) was used. Each of the reaction composed of 12.5 μl double strength master mix (Thermo Scientific, USA), 1 μl each of the primers synthesised by Inqaba Biotech (Pretoria, RSA), 5.5 μl of nuclease-free water (Thermo Scientific, USA) and DNA template in a total reaction volume of 25 μl. Negative controls were used in all reactions, which consisted of all the reaction mixture above with the exception that the DNA template was replaced with PCR buffer and nuclease-free water. Each amplified DNA (5 μl) was loaded in a 1.5% (w/v) horizontal agarose gel (Separations, South Africa) containing Ethidium Bromide (0.001 μg/ml). A DNA ladder of either 50 or 100-bp (Thermo Scientific, USA) was also added to each gel during electrophoresis to estimate the appropriate expected band size for each of the genes assayed for depending on availability. The gel run using 0.5X TBE buffer for 45 mins at 100 volts and visualised by a UV transillumination (ALLIANCE 4.7).

### Data analysis

One-way analysis of variance (ANOVA) with a 95% confidence interval was carried out to test the significance difference in the counts of the Enterobacteriaceae across the different sampling sites. The antibiotic susceptibility tests results were analysed using Microsoft Excel 2010.

## Results

### The Distribution of Enterobacteriaceae in river water

The distribution of presumptive Enterobacteriaceae in the rivers ranged from 1 × 10^2^ cfu/100ml in T2 to 1.95 × 10^4^ cfu/100ml in T4, as shown in Figure 1. The distributions of Enterobacteriaceae in rivers collected from both District Municipalities were statistically significant at *P≤* 0.05.

**Figure 1:**
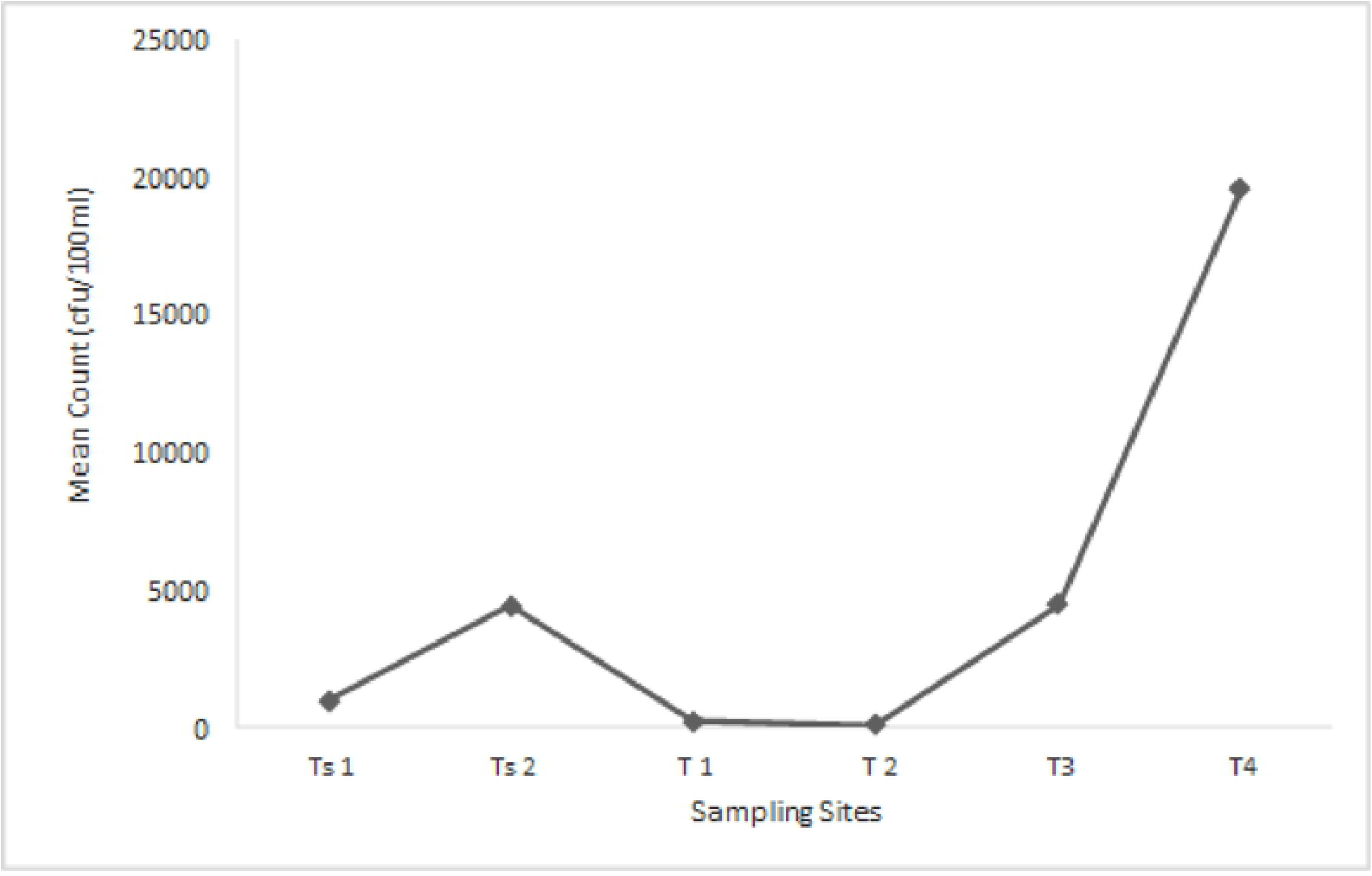
The mean of Enterobacteriaceae cell counts from Tsomo and Tyhume rivers. Tsl and Ts2 are sampling points along Tsomo river while T1 -T4 are sampling points along the Tyhume river. Using the Welch and Brown-Forsythe robust tests of equality of means, the mean counts were statistically significant at P ≤0.05 and F= 256.59. The statistics were asymptotically F distributed.

### MALDI-TOF identification of presumptive Enterobacteriaceae

Out of the 79 presumptive Enterobacteriaceae isolates recovered from the rivers, 56 were confirmed. They belonged to seven genera with species in *Citrobacter-*, *Enterobacter-*, *Escherichia-*, *Klebsiella-*, *Plesiomonas-*, *Proteus-* and *Serratia*. Other bacteria families identified include Bacillaceae (2), Morganellaceae (1), Pseudomonadaceae (5), and Staphylococcaceae (2). Thirteen of the isolates had a MALDI-TOF score value less than 1.7, and as such, their identities were not reliable and consequently excluded from this study. The full identification of the isolates showing the various genus and species identified, their numbers observed as well as the families they belong to are presented in Supplementary Table 4. The total numbers of targeted members of the Enterobacteriaceae family (*Citrobacter* species, *Enterobacter* species, *Klebsiella* species, and *Escherichia coli*) which were considered in this study was 40. Among this bacterial species, *E. coli* was dominantly isolated comprising 33% followed by *K. pneumoniae* and *E. aerogenes* which were 18% and 17% respectively of the total isolates. The distribution of the bacteria isolates is shown in Figure 2.

**Figure 2:**
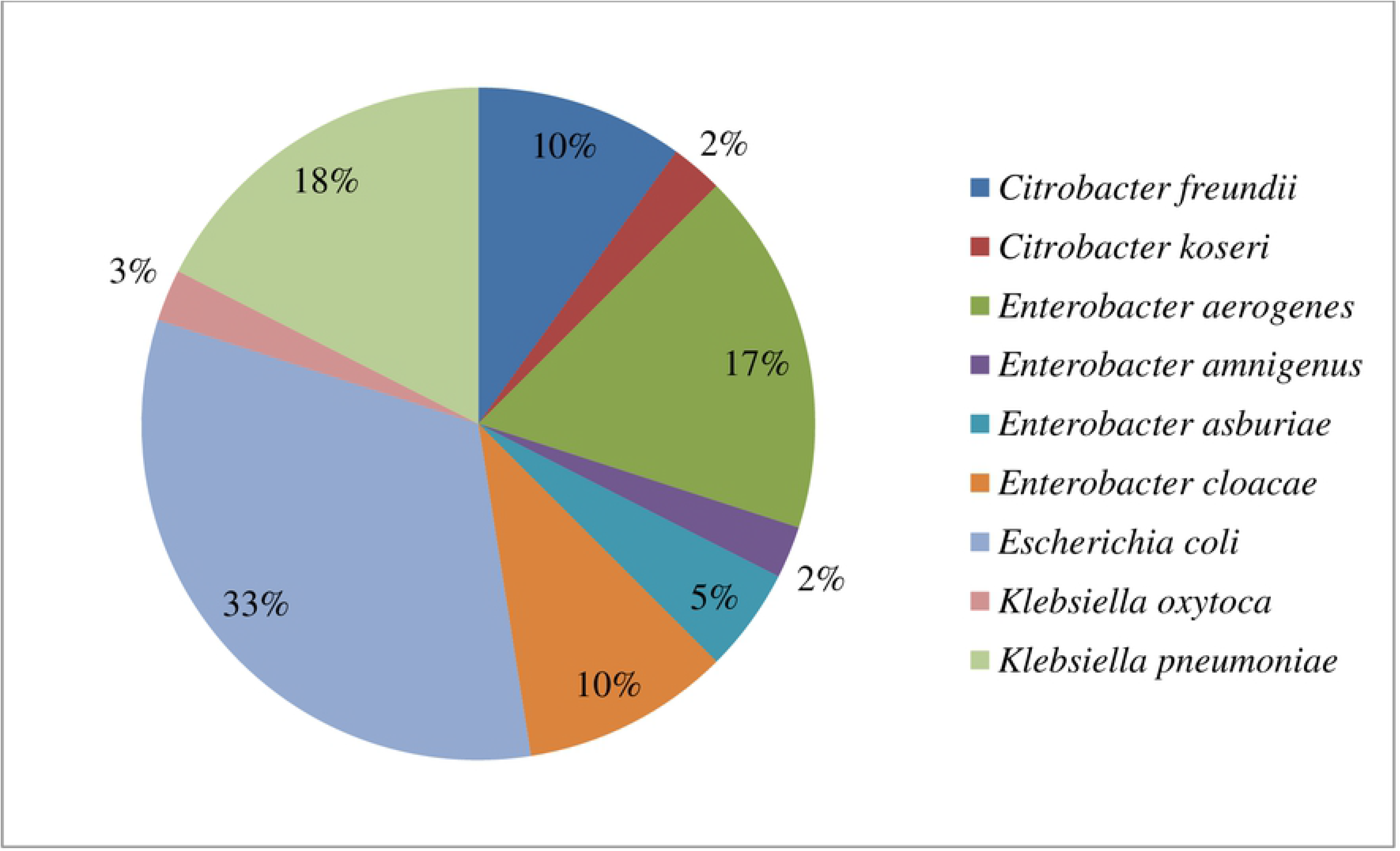
The distribution of the confirmed members of the targeted Enterobacteriaceae (n=40) recovered from the rivers.

### Evaluation of antibiotic susceptibility profile of targeted members of Enterobacteriaceae to a panel of test antibiotics

The antibiotic resistance profiles of the confirmed targeted members of Enterobacteriaceae (n=40) recovered from the rivers is shown in Figure 3. All the 14 *Enterobacter* spp. isolated exhibited resistance against the two selected antibiotics in the β-lactams (ampicillin and amoxicillin/clavulanic acid) as well as doxycycline. This was closely followed by resistance against the two polymyxins (polymyxin B and colistin sulphate) and tetracycline, where 92.9% of the isolates exhibited phenotypic resistance against these three antibiotics. The antibiotics to which the least resistance (7.1%) was observed were amikacin and imipenem. For *Klebsiella* species, all the 8 isolates exhibited resistance against ampicillin, followed by the cephems class (cefotaxime and cefuroxime) and trimethoprim-sulfamethoxazole with a resistance frequency of 87.5%. No resistance was observed against gentamicin, imipenem and meropenem amongst all the *Klebsiella* spp. from the freshwater sources. The two highest frequency of resistance observed in *E. coli* isolates to antibiotics were recorded for the class tetracyclines wherein they were all resistant against tetracycline while 92.3% exhibited resistance against doxycycline. This was followed by the resistance of 84.6% displayed across four different classes of antibiotics which include antibiotics ampicillin, nitrofurantoin, polymyxin B and trimethoprim-sulfamethoxazole. It is interesting to note that all five *Citrobacter* spp. isolated displayed resistance against at least 10 of the 18 tested antibiotics, which were displayed across 6 different classes of antibiotics. However, no resistance was observed to imipenem as all the isolates were susceptible to this carbapenem.

**Figure 3:**
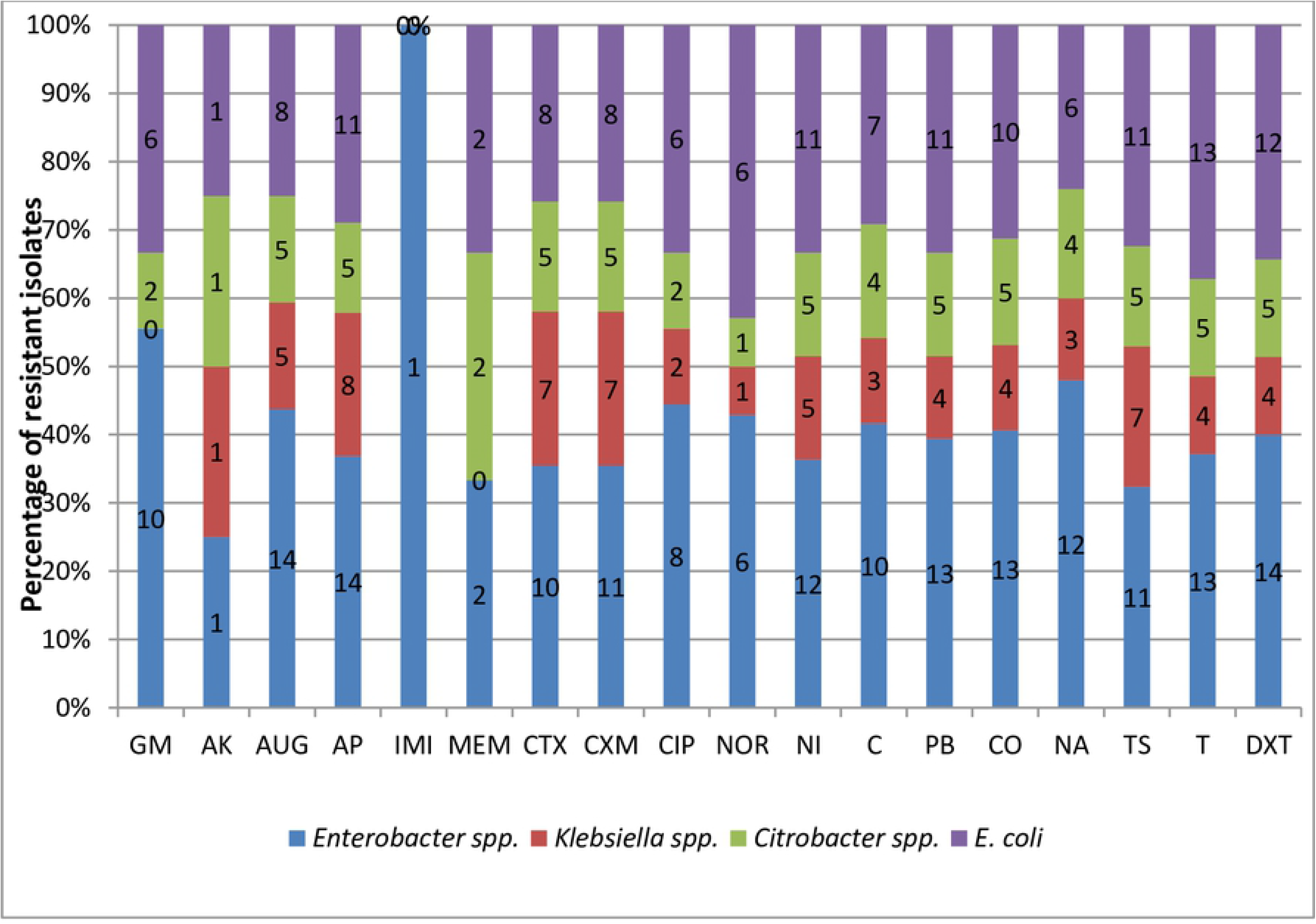
Antibiotic resistance frequencies of targeted Enterobacteriaceae recovered from Tyhume and Tsomo Rivers. *Enterobacter* spp. (n=14), *Klebsiella* spp. (n=8), *Citrobacter* spp. (n=5) and *E. coli* (n=13). Antibiotics code: GM-Gentamicin, AK-Amikacin, AUG - Amoxicillin/Clavulanic acid, AM- Ampicillin, IMI- Imipenem, MEM- Meropenem, CTX- Cefotaxime, CXM-Cefuroxime CIP-Ciprofloxacin, NOR-Norfloxacin, NI-Nitrofurantoin, C- Chloramphenicol, PB- Polymyxin B, CO-Colistin sulphate, TS-Trimethoprim/Sulfamethoxazole, T- Tetracycline and DXT-Doxycycline.

### Phenotypic antibiotic resistance pattern of targeted members of Enterobacteriaceae from rivers

Generally, varying phenotypic resistance pattern was recorded for most of the targeted members of Enterobacteriaceae recovered. The antibiotic susceptibility pattern for each bacteria isolates to each tested antibiotics is shown in the heatmap in Figure 4. The result observed indicates the effectiveness of the antibiotics towards each isolate which is an indication of the probable genetic characteristics possessed by each bacteria species. From this result, all the isolates showed full resistance against ampicillin except for isolates 32 and 33, which showed intermediate resistance. One *C. freundii* (isolate No. 24) displayed resistance against all the test antibiotics except for imipenem.

**Fig 4:**
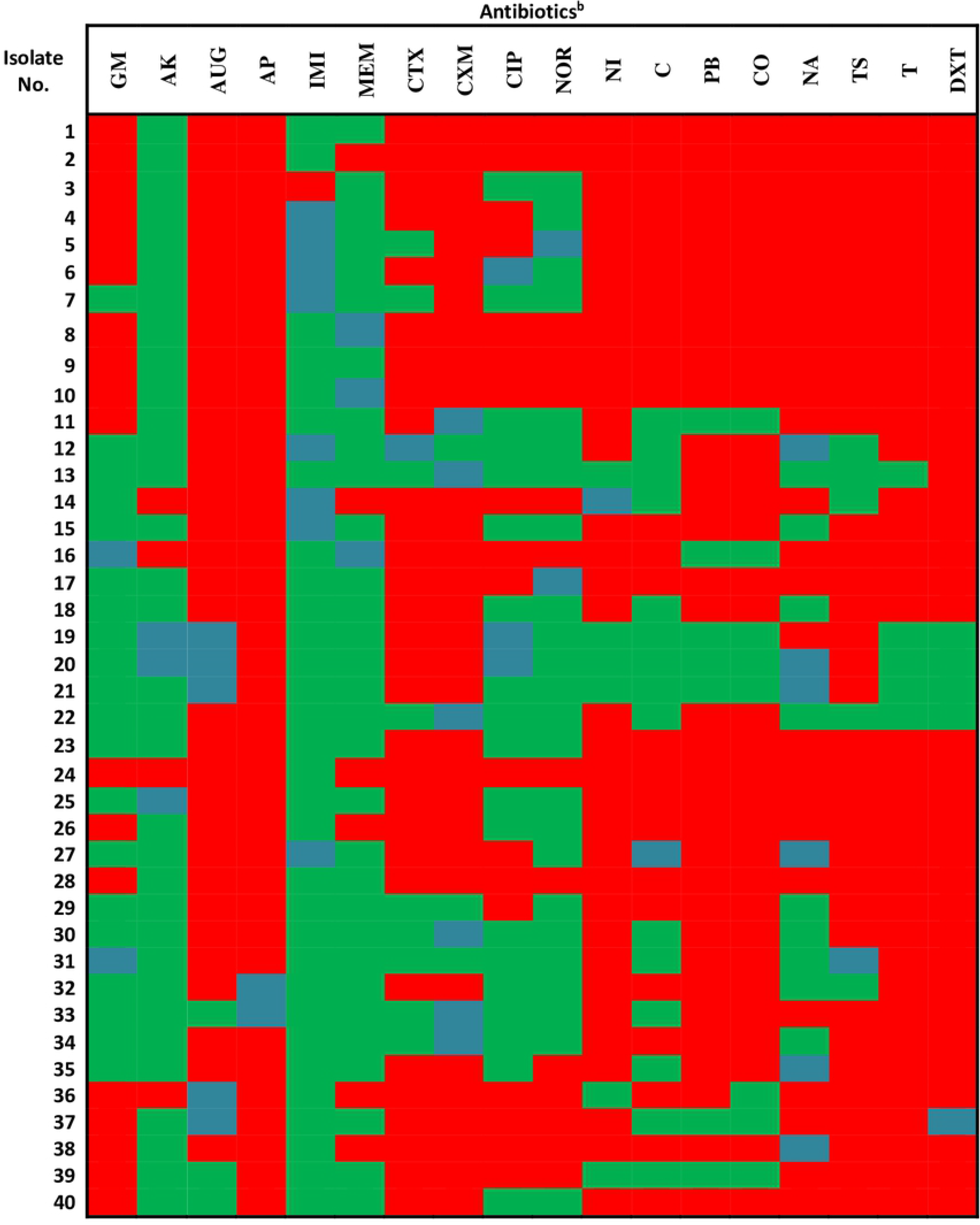
A heatmap for the antibiotic resistance patterns for each of the bacteria recovered. Colour code: Resistance (Red), Intermediate (Aqua), Sensitive (Green) Isolate No. 1 is *E. amnigenus*, 2-5 is *E. cloacae*, 6&7are *E. asburiae*, 8-14 is *E. aerogenes*, 12-21 is *K. pnuemoniae*, 22 is *K. oxytoca*, 23 is *C. koseri*, 24-27 is *C.freundii*, and 28-40 is *E. coli*. ^b^ represents anibiotics code which are exolained in Figure 3 footnotes.

### Assessment of MARP and MARI of targeted members of the Enterobacteriaceae group

The MARP patterns with the MARI exhibited by *Enterobacter* spp. and *Klebsiella* spp. are shown in Table 1 while those of *Citrobacter* spp. and *E. coli* are shown in Table 2. All the targeted bacteria recovered displayed resistance against at least four of the test antibiotics while the highest resistance pattern was observed in a *C. freundii* which was resistant against 17 out of the 18 tested antibiotics. The MARI generally ranged from 0.22 to 0.94 amongst all the isolates, which are higher than the permissible MARI value of 0.2. Majority of the phenotypes were observed uniquely.

**Table 1:**
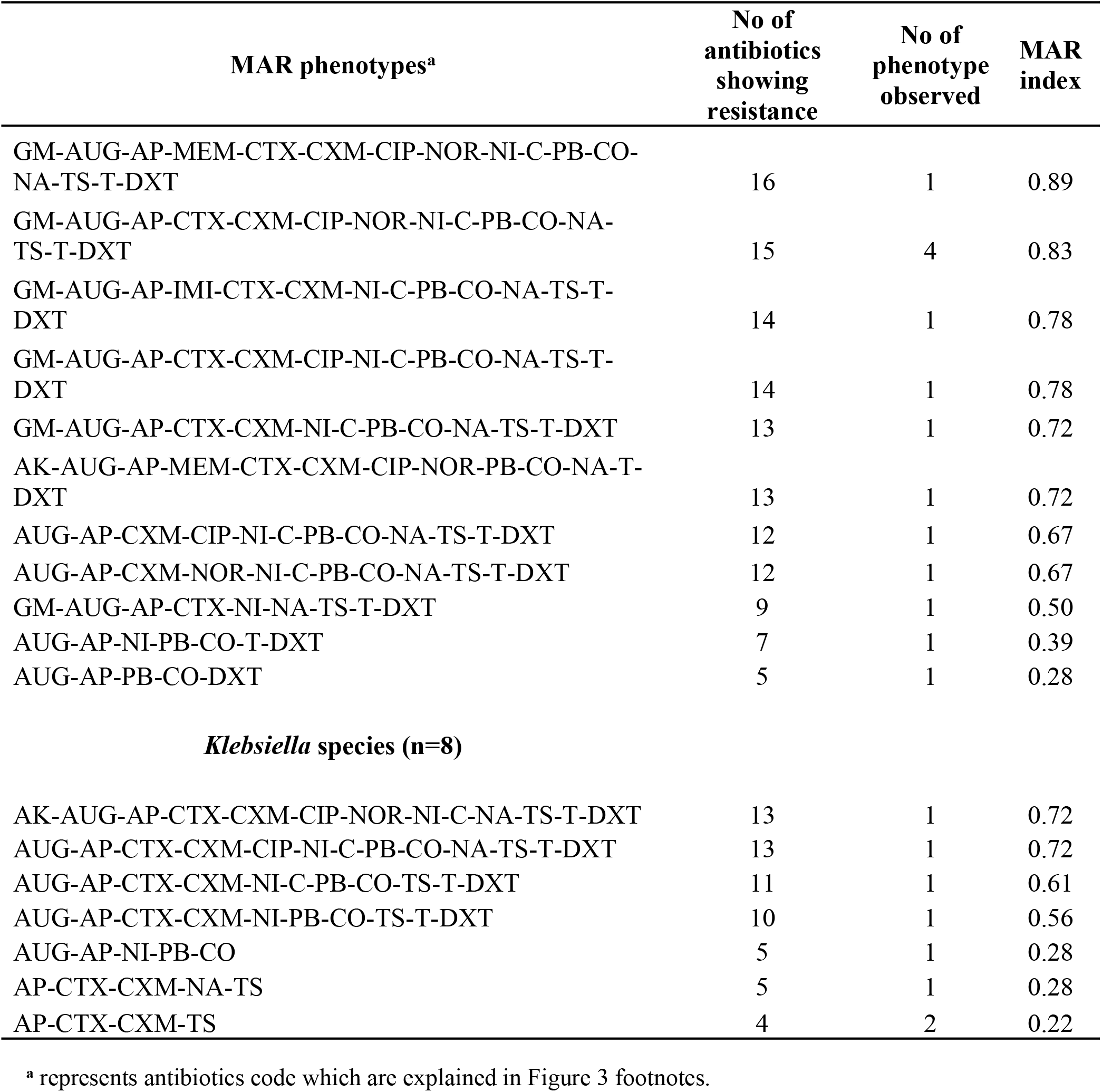
MAR phenotypes and MAR indices patterns in *Enterobacter* spp. and *Klebsiella* spp. isolated from the rivers.

| MAR phenotypes<sup>a</sup> | No of antibiotics showing resistance | No of phenotype observed | MAR index |
| --- | --- | --- | --- |
| GM-AUG-AP-MEM-CTX-CXM-CIP-NOR-NI-C-PB-CO-NA-TS-T-DXT | 16 | 1 | 0.89 |
| GM-AUG-AP-CTX-CXM-CIP-NOR-NI-C-PB-CO-NA-TS-T-DXT | 15 | 4 | 0.83 |
| GM-AUG-AP-IMI-CTX-CXM-NI-C-PB-CO-NA-TS-T-DXT | 14 | 1 | 0.78 |
| GM-AUG-AP-CTX-CXM-CIP-NI-C-PB-CO-NA-TS-T-DXT | 14 | 1 | 0.78 |
| GM-AUG-AP-CTX-CXM-NI-C-PB-CO-NA-TS-T-DXT | 13 | 1 | 0.72 |
| AK-AUG-AP-MEM-CTX-CXM-CIP-NOR-PB-CO-NA-T-DXT | 13 | 1 | 0.72 |
| AUG-AP-CXM-CIP-NI-C-PB-CO-NA-TS-T-DXT | 12 | 1 | 0.67 |
| AUG-AP-CXM-NOR-NI-C-PB-CO-NA-TS-T-DXT | 12 | 1 | 0.67 |
| GM-AUG-AP-CTX-NI-NA-TS-T-DXT | 9 | 1 | 0.50 |
| AUG-AP-NI-PB-CO-T-DXT | 7 | 1 | 0.39 |
| AUG-AP-PB-CO-DXT | 5 | 1 | 0.28 |
| *Klebsiella* species (n=8) |  |  |  |
| AK-AUG-AP-CTX-CXM-CIP-NOR-NI-C-NA-TS-T-DXT | 13 | 1 | 0.72 |
| AUG-AP-CTX-CXM-CIP-NI-C-PB-CO-NA-TS-T-DXT | 13 | 1 | 0.72 |
| AUG-AP-CTX-CXM-NI-C-PB-CO-TS-T-DXT | 11 | 1 | 0.61 |
| AUG-AP-CTX-CXM-NI-PB-CO-TS-T-DXT | 10 | 1 | 0.56 |
| AUG-AP-NI-PB-CO | 5 | 1 | 0.28 |
| AP-CTX-CXM-NA-TS | 5 | 1 | 0.28 |
| AP-CTX-CXM-TS | 4 | 2 | 0.22 |

<sup>a</sup> represents antibiotics code which are explained in Figure 3 footnotes.

**Table 2:** MAR phenotypes and MAR indices patterns in *Citrobacter* spp. and *E. coli* isolated from the rivers.

| *Citrobacter* species (n=5) |  |  |  |
| --- | --- | --- | --- |
| MAR phenotypes<sup>a</sup> | No of antibiotics showing resistance | No of phenotype observed | MAR index |
| GM-AK-AUG-AP-MEM-CTX-CXM-CIP-NOR-NI-C-PB-CO-NA-TS-T-DXT | 17 | 1 | 0.94 |
| GM-AUG-AP-MEM-CTX-CXM-NI-C-PB-CO-NA-TS-T-DXT | 14 | 1 | 0.78 |
| AUG-AP-CTX-CXM-NI-C-PB-CO-NA-TS-T-DXT | 12 | 2 | 0.67 |
| AUG-AP-CTX-CXM-CIP-NI-PB-CO-TS-T-DXT | 11 | 1 | 0.61 |
| *E. coli* (n=13) |  |  |  |
| GM-AUG-AP-MEM-CTX-CXM-CIP-NOR-NI-C-PB-CO-TS-T-DXT | 15 | 1 | 0.83 |
| GM-AUG-AP-CTX-CXM-CIP-NOR-NI-C-PB-CO-NA-TS-T-DXT | 15 | 1 | 0.83 |
| GM-AK-AP-MEM-CTX-CXM-CIP-NOR-C-PB-NA-TS-T-DXT | 14 | 1 | 0.78 |
| GM-AP-CTX-CXM-NI-C-PB-CO-NA-TS-T-DXT | 12 | 1 | 0.67 |
| AUG-AP-CTX-CXM-NOR-NI-PB-CO-TS-T-DXT | 11 | 1 | 0.61 |
| GM-AP-CTX-CXM-CIP-NOR-NI-NA-TS-T | 10 | 1 | 0.56 |
| GM-AP-CTX-CXM-CIP-NOR-NA-TS-T-DXT | 10 | 1 | 0.56 |
| AUG-AP-CIP-NI-C-PB-CO-TS-T-DXT | 10 | 1 | 0.56 |
| AUG-AP-NI-C-PB-CO-TS-T-DXT | 9 | 1 | 0.50 |
| AUG-CTX-CXM-NI-C-PB-CO-T-DXT | 9 | 1 | 0.50 |
| AUG-AP-NI-PB-CO-TS-T-DXT | 8 | 1 | 0.44 |
| AUG-AP-NI-PB-CO-T-DXT | 7 | 1 | 0.39 |
| NI-PB-CO-NA-TS-T-DXT | 7 | 1 | 0.39 |

<sup>a</sup> represents antibiotics code which are explained in Figure 3 footnotes.

### The assortment of ARGs in the Enterobacteriaceae species

A variety of genes which have been implicated in antimicrobial resistance were investigated in this study. These include the three major classes of the antibiotics; the β-lactams (amoxicillin/clavulanic acid and ampicillin); the carbapenems (imipenem and meropenem); and the cephems (cefotaxime and cefuroxime). Also investigated in this study are some other equally essential resistance genes which do not belong to the β-lactams. The genes include some resistance determinants for antibiotic classes sulphonamide, tetracycline, phenicol and aminoglycoside. Almost all of the 19 β-lactamase resistance genes assayed for were recovered in the members of the Enterobacteriaceae studied except one ESBL (*bla*_VEB_), and two pAmpC (*bla*_DHA_ and *bla*_MOX_) genes which were not detected. The frequencies of detection across the bacterial isolates are as presented in Figure 5. Also, among the 12 non-β- lactamase resistance genes assayed for, the resistances only *tetK* and *StrA gene* were not detected in this study. These represented the tetracyclines and aminoglycosides, respectively. The frequencies of detection of these genes are also depicted in Figure 6.

**Figure 5:**
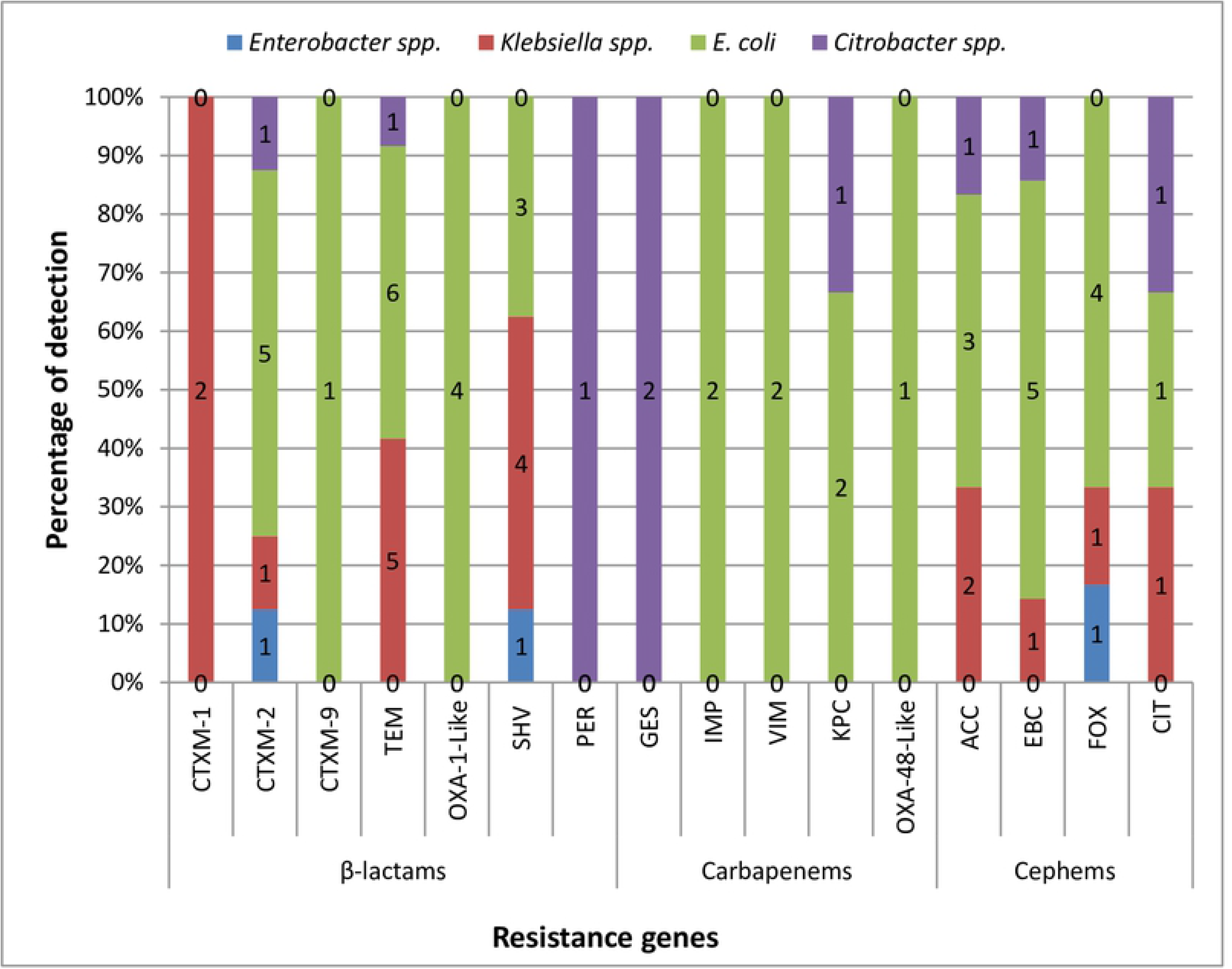
The prcvalcncc of key β-lactamases genes detected across the bacterial isolates. These include the pAmpC genes coding for resistance against the cephems class of antibiotics and other ESBL resistance encoding genes. For β-lactams and Cephems: *Enterobacter* spp. (n=14), *Klebsiella* spp. (n=8) and *Citrobacter* spp. (n=5); β-lactams: *E. coli* (n= 13); Cephems: *E. coli* (n=l 1); Carbapenems: *E. coli* (n=2) and *Citrobacter* spp. (n=3), where n is the number of phenotypic resistant isolates screened for resistance genes.

**Figure 6:**
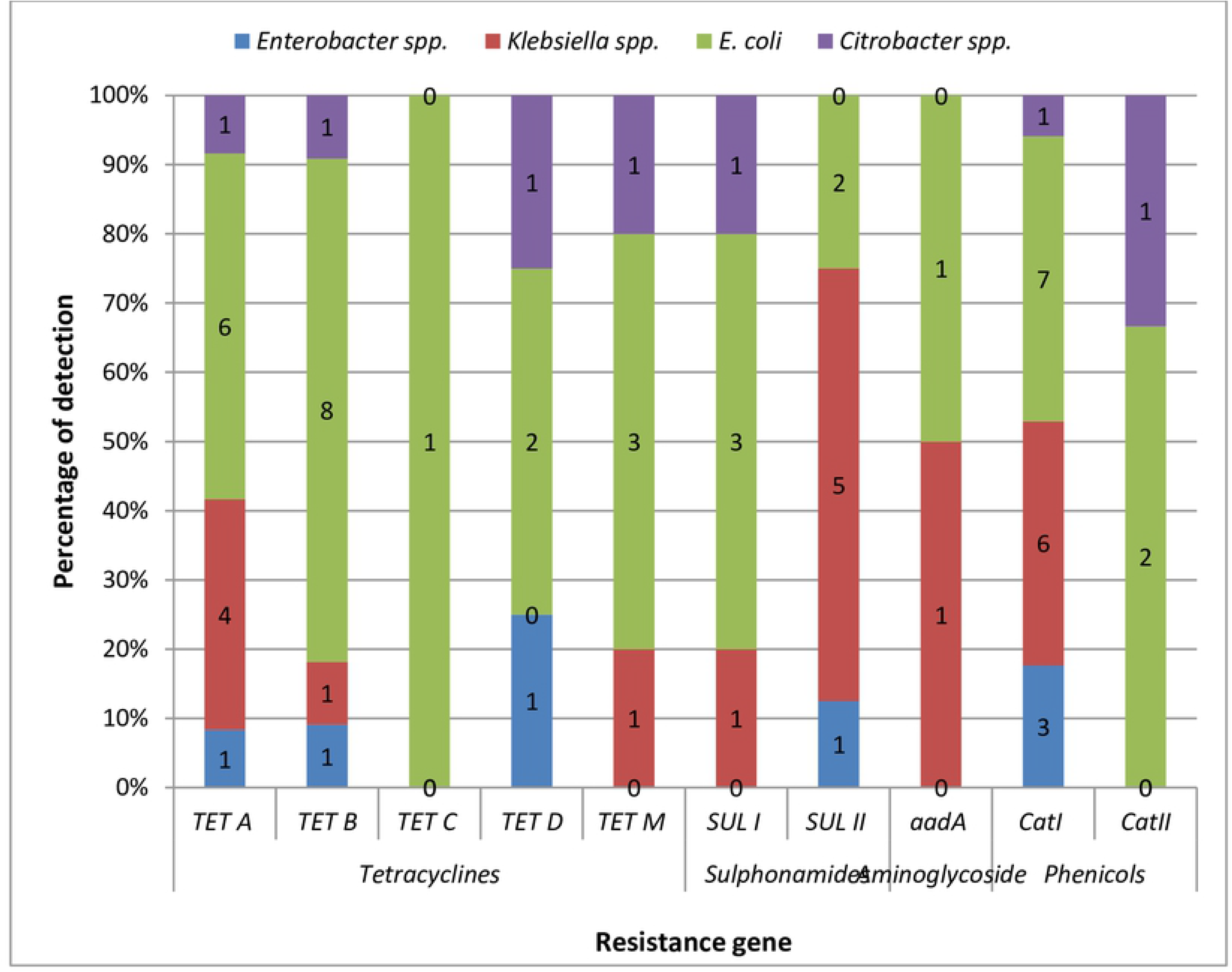
The prevalence of key non-β-lactamase resistance genes detected across the bacterial isolates. These include the genes coding for resistance against antibiotic classes tetracycline, sulphonamide, aminoglycoside and phenicol. For tetracyclines, sulphonamides, aminoglycosides and phenicols: *Enterobacter* spp. (n=14, 11, 11 and 10 respectively), *Klebsiella* spp. (n=4, 7, 3 and 3 respectively), *E. coli* (n=13, 12, 7 and 7 respectively); *Citrobacter* spp. (n=5, 5, 3 and 5 respectively) where n is the number of phenotypically resistant isolates screened for resistance genes.

For the β-lactamase resistance determinants, all the isolates harboured the *bla*_TEM_ and *bla*_SHV_ genes with exception to *Enterobacter* spp. and *Citrobacter* spp. respectively. The most frequently detected *bla*_TEM_ was observed in *E. coli* (50%) while the isolates harbouring most of the *bla*_SHV_ was among the *Klebsiella* spp. (50%). The OXA-1-like gene was only detected in *E. coli* while only one *Citrobacter koseri* harboured *bla*_PER_. At least one of the CTX-M genes (group1, 2, 9) was detected across all the isolates. The majority of the carbapenemase resistance determinants were harboured by *E. coli*. *Bla*_IMP_, *bla*_VIM_ and *bla*_OXA-48-like_ ARGs were found only in *E. coli* while *bla*_GES_ was only harboured by *Citrobacter* spp. The *bla*_KPC_ gene was harboured in *E. coli* and *Citrobacter* spp. For pAmpC, the resistance determinants *bla*_ACC_, *bla*_EBC_, *bla*_FOX_ and *bla*_CIT_ genes were all harboured by the isolates except *bla*_ACC_ and *bla*_FOX_ which were not detected in *Enterobacter* spp. and *Citrobacter* spp. respectively.

Among the non-β-lactamase resistance genes assayed for, the most frequently detected was the *catII* encoding resistance to the class phenicol. All of the *E. coli* and *Klebsiella* spp. recovered harboured this *catII* gene. Meanwhile, other gene assayed for in the phenicol resistant isolates, *catI* genes, were harboured only in *E. coli* and *Citrobacter* spp. isolated with 67% and 33% frequencies respectively. In the class aminoglycosides, *aadA* gene was harboured only in *Klebsiella* spp. (50%) and *E. coli* (50%) isolated. Among the genes assayed for tetracycline resistance, the highly detected was *tetA* followed by *tetB*, *tetM*, and *tetD, tetC* gene in that descending order. The *tetC* gene was the least detected, which was harboured by only one *E. coli* isolate. For the sulphonamides, the gene *sulII* was more detected with 8 of the isolates harbouring this gene than its *sulI* counterpart which had only 5 of the isolates harbouring the gene. The *aadA* aminoglycoside gene was detected at an equal rate in *Klebsiella* spp. and *E. coli* while the *StrA* gene was not harboured at all by any of the targeted Enterobacteriaceae in this study.

Various detected genotypic patterns of the MDR Enterobacteriaceae were also described in Table 3. The ESBLs genes detected included *bla*_CTX-M_ (including group 1, 2 and 9), *bla*_TEM_, *bla*_VIM_, *bla*_OXA-1-like_, *bla*_OXA-48-Like_, *bla*_SHV_, *bla*_PER_, *bla*_GES_, *bla*_IMP_ and *bla*_KPC_. The pAmpC genes which were detected include *bla*_ACC_, *bla*_EBC_, *bla*_FOX_ and *bla*_CIT_. The non-β-lactam resistance genes detected include *tetA*, *tetB*, *tetC*, *tetD*. *tetM*, *sulI*, *sulII, aadA*, *catI* and *catII.* Out of the 40 MDR Enterobacteriaceae strains, 28 harboured at least one of the resistance genes assayed. Notably, all the genotypic patterns occurred uniquely.

**Table 3:**
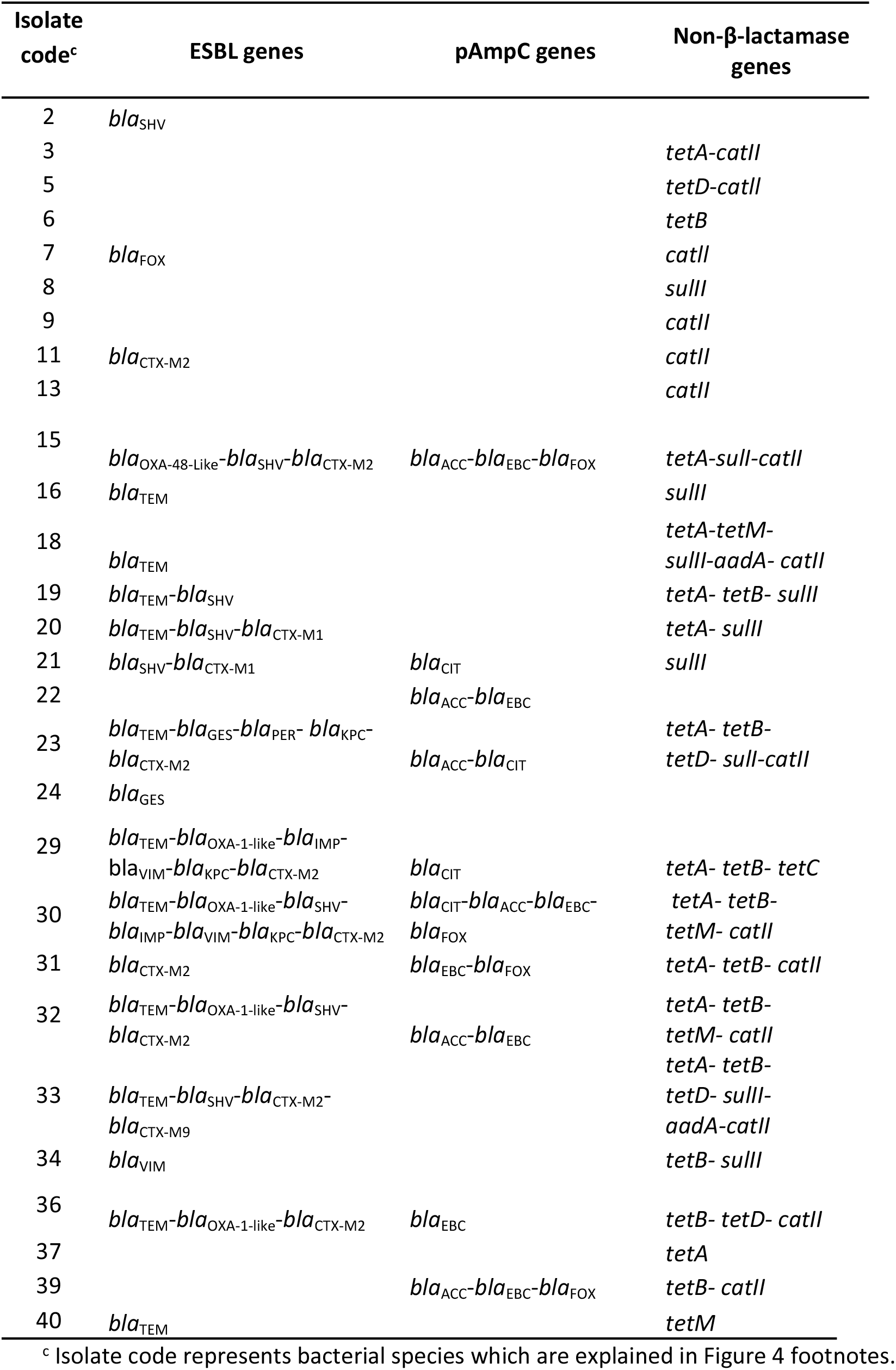
Extended-spectrum and plasmid-mediated AmpC β-lactamases, and non-β-lactamase genotypes detected in multidrug-resistant Enterobacteriaceae strains.

## DISCUSSION

The distribution of presumptive Enterobacteriaceae in the rivers sampled in this study shows that coliform bacteria are widely distributed in aquatic environments. The count range observed fall within the limit of between 100 and 100,000 CFU/ 100ml for polluted surface water by (52). A study by (53) on bacteriological qualities of Tyhume river suggested that the quality of the river water is poor and this corroborates the finding in this study that coliform bacteria impair the water quality. The benchmark value for faecal coliforms being 0 CFU/100ml is set in South Africa, for domestic water (54) and this was employed for the interpretation of the data obtained. This is necessary because, across all sampling points, the water is being used for a variety of domestic purposes. The detection and subsequent confirmation of Enterobacteriaceae from the rivers may be attributable to the probable point and non-point pollution across the various sampling points. Observations revealed that the rivers at the sampling sites also serve as sources of drinking water for livestock and irrigation, among other purposes such as recreational (swimming) and religious (baptism) activities. It is also important to note that, upstream of the sampled points, the rivers received wastewater runoffs from surrounding communities. These activities highlighted could contribute substances that are capable of inducing resistance in microorganisms either directly or indirectly and could be responsible for the abundance of Enterobacteriaceae in these water sources. Surface water sources are considered to be reservoirs of microorganisms, metals and antibiotic resistance genes. Thereby plays a vital role in not only the spread but also in their active transport since surface water sources are recipients of wastewaters harbouring contaminants from diverse sources (5,55–57).

The peaks of the presumptive Enterobacteriaceae were observed at T4 and TS2 for Tyhume and Tsomo rivers respectively. It is highly pertinent to state that these sites with the highest counts are the only ones with semi-urban settlements on the different river courses. There is a probable higher pressure of anthropogenic activities around these sites necessitating a higher than usual bacterial counts. The highest distribution of the presumptive Enterobacteriaceae observed at point T4 in Alice along the course of the Tyhume river could not be unconnected with the discharge from a municipal WWTP upstream and higher pressure of anthropogenic activities observed around the area during sampling. This higher than usual bacterial count obtained is relatively close to what was described by (53). Most research concepts have employed methods that allow for the enumeration of total bacteria count irrespective of the prevailing conditions during sampling. Some studies focused on the enumeration of selected members of the Enterobacteriaceae group rather than assessing the collection of species in the different aquatic environment. The different approaches have however indicated that the Enterobacteriaceae group had become a common feature in much surface water globally. The presence of faecal pathogens could have a grave public health concern, especially in developing countries where dependence on surface water sources is still high due to inaccessibility to potable water for domestic and other uses. Bringing into perspective the water quality guideline, the quality of the rivers across all the sites in this study were below the standard limits, thereby making them unsuitable for domestic utilisations.

The MALDI-TOF confirmation of the identities of the presumptive Enterobacteriaceae isolated revealed that *E. coli* was mostly detected in this study with a 33% detection rate. A similar result was reported by (10) and (11). This high detection rate of *E. coli* when considered with the other members of Enterobacteriaceae detected is, however, unsurprising as *E. coli* is a common commensal organism in the gastrointestinal tracts of animals as well as humans. In developing countries, one of the major causes of waterborne disease outbreaks is due to the consumption of contaminated water which significantly harbours gastrointestinal microbial pathogens. Rivers are generally considered as reservoirs of ARB more mainly because they are recipients of surface waters containing materials from different origins.

These sources include effluents from WWTPs, urban and industrial effluents which harbour an immense antibiotic resistome comprising of both pathogenic and non-pathogenic bacteria (38,58). The presence of ARB in rivers is a significant public health concern. This is because they can be transferred to humans when the contaminated water are consumed either directly as drinking water, or indirectly during other uses such as during recreational activities, religious activities, abstracted for irrigation, just to mention a few. The use of these rivers, in turn, can contribute to the rapid and aggravated spread and persistence of ARB within the general populace and environment. Effluents of low quality from WWTPs, hospitals and abattoirs can easily contaminate the water-bodies that receive this discharge with potential MDR pathogenic Enterobacteriaceae species. Although this current study did not extend its investigation into the pathogenic potential of the bacterial isolates, however, all the selected Enterobacteriaceae isolates revealed a widespread presence of MDR bacteria in Tyhume and Tsomo Rivers.

Owing to the phenotypic antibiotic resistance pattern obtained, all the selected Enterobacteriaceae isolates recovered from these rivers exhibited multiple antimicrobial resistance patterns with the lowest resistance pattern being against four antibiotics and the highest resistance pattern being against seventeen of the eighteen tested antibiotics. Majority of these resistance patterns occurred singly with three exceptions. These include a MAR phenotype pattern among *Enterobacter* spp. with resistance to fifteen antibiotics that occurred 4 times, a pattern among *Klebsiella* spp. and *Citrobacter* spp. each occurred twice. The breakdown of the patterns of the targeted Enterobacteriaceae isolates (40) in these rivers are as follow; four-antibiotics (2), five-antibiotics (3), seven-antibiotics (3), eight-antibiotics (1), nine-antibiotics (3), ten-antibiotics (4), eleven-antibiotics (3), twelve-antibiotics (5), thirteen-antibiotics (4), fourteen-antibiotics (4), fifteen-antibiotics (6), sixteen-antibiotics (1) and seventeen-antibiotics (1) as seen in Table 1 and 2. This antimicrobial resistance pattern indicates that all the isolates displayed resistance against multiple antibiotics. The mean MAR index was 0.61, and the value ranged from 0.22 to 0.94, indicating heavy contamination of the rivers probably with wastewaters. The occurrence of MARPs in bacteria isolates of enteric origin found in aquatic environment has been previously documented (9,59–61). In KwaZulu-Natal Province, RSA, 71.15 – 97.1% of *E. coli* isolated from two rivers in Durban exhibited MDR against the antibiotics tested (59). Titilawo and colleagues (9) also reported multiple antimicrobial resistances ranging from three to nine antimicrobials from *E. coli* isolated from some rivers in Nigeria. Toroglu and colleagues (62) reported MDR in the members of Enterobacteriaceae recovered from some rivers in Turkey (62) with similar results being reported by (63) and (11) in the rivers of Bangladesh and Ethiopia respectively. Various human activities contributes to the diverse MARPs observed in rivers. One of the most important being the discharge of wastewaters as rivers are the primary recipients of effluents from sewage (11,64). It is rather unfortunate that majority of the antimicrobial agents when used are not fully metabolised in the body and are therefore released directly into the hospital sewage system or to the municipal wastewater (5,23). Final effluents of the municipal wastewater are in turn discharged directly into surface water. Many studies have emphasised that polluted environmental sources are mostly responsible for the promotion of ARGs (8,12,26). The MARI of all the isolates in this study was higher than 0.2, which is an indication that these isolates were culled from high-risk environments where there is a high selective pressure of antibiotic resistance. The results obtained from this study, in general, is very close to those obtained by (65) and (66) from clinical settings wherein their isolates also exhibited high level of resistances against β-lactams, with the exception to carbapenems which are usually the last resort antibiotics for therapy against infections caused by MDR ESBL-producing Enterobacteriaceae. Likewise, a similar trend was reported by (67) for Enterobacteriaceae recovered from rivers. The aquatic environment thus remains a vital pathway/ reservoir for the spread of antimicrobial resistance in the environment.

Members of Enterobacteriaceae which are natural commensals of humans and animals and those often implicated in nosocomial and community-acquired opportunistic pathogens play a significant function in the movement of ARGs in microorganisms of environmental origin to other species and finally reaching human which can then be pathogenic (26,68). This course has been tackled by the epidemiology of the dissemination of ESBL/pAmpC in aquatic milieu (26,69–71). Here, the genetic characterisation of essential *bla* genes (ESBLs and pAmpC) among β-lactam-resistant Enterobacteriaceae, which include the carbapenem-resistant Enterobacteriaceae as well as the cephem-resistant Enterobacteriaceae revealed a high occurrence rate of various resistance determinants. The results obtained showed that *E. coli* and *Klebsiella* isolates harboured more of these resistance genes assayed. The higher rate of resistance observed in these two genera of Enterobacteriaceae studied could be due to their propensity to undergo stable acquisition and integration of ARGs in the aquatic environment than other species recovered. We observed twenty-eight different resistance genotype patterns in the MDR isolates, indicating that the isolates have independent clusters of genes responsible for their resistance phenotypes. The ease of the acquisition of ARGs in *E. coli* have been reported (40,72). In the context of carriage of β-lactamase genes, 87.5% of *Klebsiella* spp. and 69% of *E. coli* recovered harboured at least one of the genes. Based on this study, pAmpC-producing were harboured in 46% of the *E. coli* isolated while only 2.8% was detected among the *Klebsiella* spp. A variety of ESBLs were also detected. The CTX-M-2 was more predominant among the CTX-M allelic variants and was harboured mostly in *E. coli* recovered. Per this study, the CTX-M-2 variant is one of the most prevalent in South Africa (73,74). In contrast, CTX-M subtypes (groups 1,2 and 9) reported in this study have also been reported in similar studies to be the most prevalent ESBLs in *E. coli* isolated from aquatic and clinical settings (75–78), human faecal isolates (79) and sewage sludge (80). However, in this study, the *bla*_TEM_ genes were the overall predominant ESBL group. The dominance of bla_TEM_ could be as a result of their earlier emergence than other β-lactamases and an indication that the selective pull favouring the ESBL genes are present in aquatic environments in South Africa. In support of our findings, the dominance of *bla*_TEM_ in *E. coli* has been reported in aquatic environments (9,81). The *bla*_CTX-M_ genes (which include the group 1, 2, and 9) which keenly followed highlights the widespread of these genes in hospital and community settings in the last decade (82). In some studies, the dominance of CTX-M genes in many different surface water has been reported (39,83,84).

It is important to highlight that simultaneous occurrence of ESBLs and pAmpCs was seen in 20% of the overall isolates recovered in this study. More importantly, co-carriage of the ESBL/pAmpC genes was most prevalent in *E. coli* with 75% occurrence. The co-occurrence of ESBL/pAmpC in *E. coli* correlated conspicuously with the result of another study which investigated the distribution of β- lactamases genes in irrigation water in South Africa (73). The most frequently detected genes was observed in one *E. coli* isolate, which harboured thirteen of the genes which included five ESBLs, four pAmpCs and four non-β-lactamase genes. However, notably among the isolates was a *C. koseri* which harboured twelve out of the genes assayed. The resistance genes detected include five of the ESBL genes (including the only *bla*_PER_ detected in this study), two of the pAmpC resistance genes and five of the non-β-lactamase genes. To the best of our knowledge, this study provides the first report of *C. koseri*, exhibiting co-occurrence of ESBL/pAmpC in environmental isolates. The detection of a *bla*_PER_ producing *Citrobacter* spp. in this study is impressive. This emerging ESBL gene has not been previously reported in environmental isolates in South Africa. Although *bla*_PER_ ESBL genes has been reported in *Aeromonas* from European surface waters and Austrian activated sludge, however this enzyme is rarely reported in clinical isolates around the world (85–87). It is interesting to note that Carbapenemase-Producing Enterobacteriaceae (CPE) isolates *bla*_KPC_, *bla*_GES_, *bla*_OXA-48-like_ and other variants *bla*_VIM_, *bla*_IMP_ that are often detected in clinical settings were also recovered in this present study. The *bla*_KPC_, *bla*_GES_ and *bla_O_*_XA-48-like_ have been found in clinical settings in South Africa (88). Although, it is unclear where these CPE isolates have emerged from whether from humans or hospital settings or municipal discharges, what remains apparent is that these isolates may spread from the aquatic environment to clinical settings in the most foreseeable future. The detection of carbapenemase and ESBL genes in rivers is worrisome and presents an indication that the dissemination of resistant strains to the environment is currently going on in South Africa.

A high prevalence of non-β-lactamases was also detected in this study. This include *tetA* (33.3%), *tetB* (30.5%)*, tetM* (13.9%), *tetD* (11.1%), *tetC* (2.8%), *catII* (68%), *catI* (12%), *sulII* (22.8%), *sulI* (14.3%), and *aadA* (8.3%). Most of these isolates recovered from these rivers harboured multiple antibiotic resistance genes which include the ESBLs, pAmpC, as well as the non-β-lactamases. The combination of these genes in MDR Enterobacteriaceae can potentially result in disastrous effects when transferred into humans either directly while utilising these water sources for various domestic purposes or indirectly when the water is abstracted for irrigation of fresh produce which usually require minimal or no processing before consumption.

## CONCLUSION

The findings from this study exposed the high occurrence of MDR Enterobacteriaceae in Tsomo and Tyhume rivers, thus requiring special attention so as not to compromise the water-food-public health interphase. The high frequency of detection of the various ARGs investigated is an indication that the MARP observed is mostly due to these genes that confer resistance against various antibiotics covered in this study. ARGs that are widely found in Enterobacteriaceae have been established to be harboured on various mobile genetic elements that play a critical function in the rapid spread of ARG determinants. Our findings support the need for comprehensive control of ARG dissemination into the environment. This is necessary due to the serious public health concern attached to the proliferation of microbial pathogens into the environment and the menace of antibiotic resistance. This can be achieved by proper management of antibiotic use in human and animal husbandry, effective management of waste (hospital, municipal, agricultural, and industrial) containing antimicrobial resistance bacteria and ARGs and lastly the management of already contaminated environments. All of the above-mentioned strategies are important in forestalling the scourge of antimicrobial resistance spread in the environment.

